# JSP1 Regulates Neutrophil Adhesion via Integrin-SRC Signaling in Vascular Inflammation

**DOI:** 10.1101/2025.09.04.674122

**Authors:** Li Li, Nicholas K Tonks

## Abstract

The c-JUN N-terminal kinase (JNK) signaling pathway plays an important role in regulating the innate immune response. Immune signaling is governed by the coordinated activity of protein kinases counter-balanced by protein phosphatases; however, the importance of the latter family of enzymes is less well understood. c-JUN N-terminal kinase (JNK)-stimulatory phosphatase 1 (JSP1, also known as DUSP22) has been implicated as a positive regulator of JNK signaling, yet its role in innate immunity is not clear. Using a mouse model of the local Shwartzman reaction, we show that JSP1 is essential for LPS-TNFα-induced vascular injury. JSP1-deficient mice exhibited reduced vascular hemorrhage. Neutrophil depletion and adoptive transfer experiments confirmed that JSP1-expressing neutrophils mediate this injury. JSP1 was not required for neutrophil development or surface receptor abundance but was essential for integrin activation and adhesion. Reduced SYK and HCK phosphorylation in JSP1-deficient neutrophils are consistent with a mechanism involving impaired integrin-SRC signaling. These findings establish JSP1 as a key regulator of neutrophil-driven vascular inflammation.

## Introduction

Mitogen-activated protein kinase (MAPK) signaling pathways are critical regulators of cellular responses to a wide range of extracellular stimuli, including inflammatory cytokines and environmental stress. These pathways orchestrate key biological processes by relaying signals from the cell surface to the nucleus, ultimately influencing gene expression and cellular behavior (1, 2). Among the MAPK cascades, the c-JUN N-terminal kinase (JNK) pathway is particularly responsive to pro-inflammatory cytokines such as tumor necrosis factor-α (TNF-α) and lipopolysaccharide (LPS), playing a pivotal role in inflammatory and stress responses.

Given the profound impact of MAPK signaling on cellular physiology, tight regulation of MAPK activity is essential. This regulation is achieved at several levels. Activation of MAPKs is coordinated in kinase cascades, the assembly of which is regulated by scaffold proteins, and counterbalanced by the action of protein phosphatases (3, 4). Members of the protein tyrosine phosphatase (PTP) family, including dual-specificity phosphatases (DUSPs), regulate MAPKs by removing phosphate groups from both threonine and tyrosine residues within their activation loops (4-6), which suppresses MAPK function and serves to switch off signaling. Furthermore, beyond direct effects on MAPKs themselves, the activation state of MAPK signaling modules can be influenced by dephosphorylation of other critical components in the cascade as well as by the organization of cascade components into ordered complexes.

JNK-stimulatory phosphatase 1 (JSP1), also known as DUSP22, JKAP, or VHX, is a member of the DUSP family that is characterized by intrinsic phosphatase activity but lacking the cdc25-homology domain typically required for substrate docking of the MAPKs themselves (7-9). Unlike most DUSPs that inactivate MAPKs, JSP1 has been shown to act as a positive regulator of the JNK pathway (7, 8). Interestingly, JSP1, and VHY/DUSP15 (10), are unique among the members of the PTP family in being myristoylated at Gly2 in the N-terminus, a modification that is required for its ability to activate JNK signaling and apoptosis (11). Although it was also suggested initially that DUSP22 dephosphorylated ERK (9), a more recent study confirmed the ability of DUSP22 to activate JNK, suggesting also that it may serve as a scaffold to promote signaling through a ASK1-MKK7-JNK pathway (12).

A variety of substrates have now been identified for JSP1/DUSP22 that expand our understanding of its biological influence. In general terms, JSP1/DUSP22 has been associated with regulation of the Epithelial-to-Mesenchymal transition (EMT), through effects on SMAD and MAPK signaling pathways (13), as well as control of cell migration, through dephosphorylation and inactivation of focal adhesion kinase, FAK (14). Dephosphorylation of FAK also underlies the ability of JSP1 to attenuate MASH (metabolic dysfunction-associated steatotic liver disease) and HCC (hepatocellular carcinoma) (15). Overexpression of JSP1 in 293T cells reduced IL-6-induced phosphorylation of STAT3 at tyrosine 705, suggesting a negative regulatory role in IL-6/STAT3-mediated signaling (16). JSP1 also negatively regulates estrogen receptor-α (ERα)-mediated transcription by dephosphorylating ERα at serine 118 (17).

JSP1 has also been implicated it in the etiology of important disease states. Knockdown of JSP1, or pharmacological inhibition of its activity, in multiple models of muscle wasting, suppressed JNK activation and prevented muscle loss via down regulation of FOXO3a, a master transcription factor promoting muscle catabolism (18). A tumor suppressor function has been identified in lung cancer. The exosomal micro-RNA miR-1228-5p, which is upregulated in small cell lung cancer, inhibits expression of JSP1 thereby facilitating cancer cell growth and metastasis (19). In non-small cell lung cancer (NSCLC) down-regulation of JSP1 coincides with decreased survival; it has been reported that JSP1 dephosphorylates AKT, underlying inhibitory effects on cell viability and migration (20). In lung adenocarcinoma, JSP1 suppresses tumorigenesis by inhibiting EGFR signaling (21); similar observations were reported in prostate cancer (22). In addition, JSP1 is an important immune regulator. In T cell receptor (TCR) signaling, JSP1 negatively regulates T cell activation by dephosphorylating LCK at Y394 (23) and modulates LCK stability through dephosphorylation-induced proteasome degradation of E3 ligase UBR2, an upstream activator, to switch off TCR signaling (24). Downregulation of JSP1 has also been linked to inflammatory disorders. Ablation of JSP1 promotes atherosclerosis through regulation of ERK and NF-κB pathways (25). Reduced JSP1 levels have also been observed in T cells of patients with ankylosing spondylitis (26) and systemic lupus erythematosus (27), in the serum of sepsis patients (28), and in synovial tissue and serum of rheumatoid arthritis patients (29). Similarly, JSP1 levels are decreased in patients with fatty liver disease (15). Collectively, these findings suggest that JSP1 may serve as a biomarker for inflammatory diseases and a potential indicator of disease prognosis.

Such observations underscore the cell- and context-specific functions of JSP1 in immune and metabolic signaling; however, its role in innate immune responses, particularly in vascular inflammation, remains poorly understood. To address this gap, we investigated the function of JSP1 in a mouse model of vascular inflammation, the local Shwartzman reaction (LSR). The LSR is elicited by sequential exposure to bacterial lipopolysaccharide (LPS) followed by tumor necrosis factor-α (TNFα), leading to robust activation of JNK signaling and local intravascular inflammation (30, 31). This response is characterized by neutrophil infiltration, cytokine release, endothelial activation, and vascular permeability changes, making it a powerful experimental tool for dissecting molecular mechanisms of inflammation-driven vascular injury. Our study reveals that JSP1 is essential for neutrophil-mediated vascular injury by regulating integrin-dependent adhesion through SRC family kinase signaling. These findings uncover a novel role for JSP1 in neutrophil activation and suggest its potential as a therapeutic target in inflammatory vascular diseases.

## Results

### JSP1 is required for LPS-induced vascular injury

JSP1 was originally identified and characterized as a selective activator of the MAPK JNK in cell co-transfection assays (7, 8). Members of the JNK group are predominantly activated after exposure of cells to proinflammatory cytokines. Therefore, to investigate further the physiological role of JSP1, we utilized a modified mouse model of the local Shwartzman reaction (LSR) (32). In this model, consecutive injections of LPS and TNFα activate JNK signaling, which provides an opportunity to explore the functions of JSP1 in the host response to inflammation stimuli.

Wild-type (WT) and JSP1-knockout (*Jsp1*^−^/^−^) mice were subjected to subcutaneous injections of LPS, followed by tumor necrosis factor-alpha (TNFα), to enhance the inflammatory response. Notably, *Jsp1*^−^/^−^ mice exhibited reduced inflammatory reactions compared to their WT counterparts, as illustrated by decreased vascular permeability (Figure 1). These observations suggest that JSP1 may play a crucial pro-inflammatory role in LPS-induced vascular injury, potentially by modulating signaling pathways involved in immune activation.

**Figure 1.**
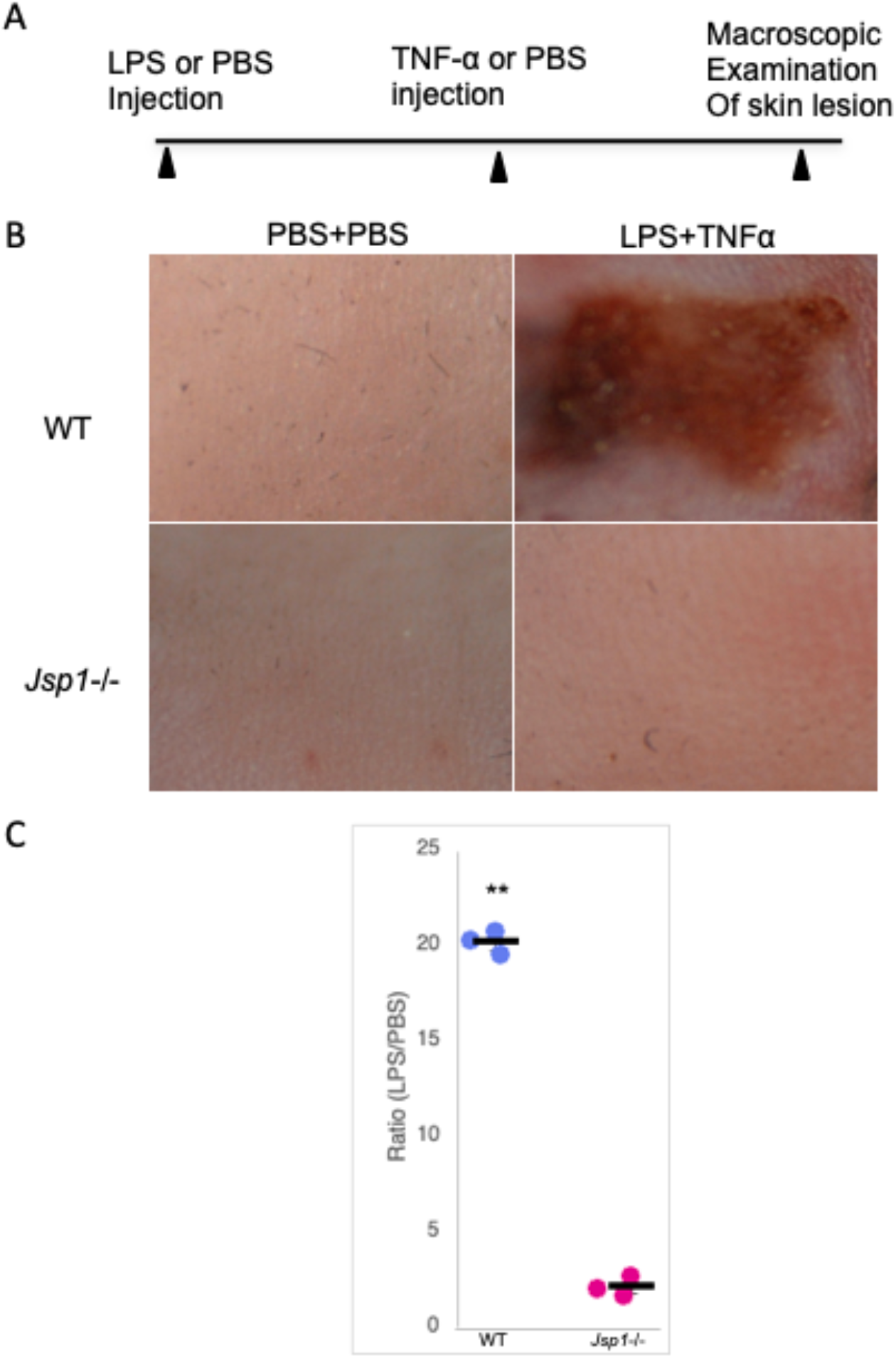
JSP1 was required for LPS-induced vascular injury. (A) Schematic representation of the injection regimen for the LSR mouse model. (B) Macroscopic appearance of dorsal skin in mice injected with PBS or LPS-TNF-α. (C) The degree of hemorrhage in the WT and *Jsp1-/-* mice was quantified by densitometry analysis of skin samples treated with either PBS or LPS injection. Each dot represents the ratio of the degree of hemorrhage obtained from a pair of mice (LPS–TNF-α–treated over PBS-treated). \*\**P* < 0.01, based on three pairs of mice per group.

### JSP1-dependent neutrophils drive LPS-TNFα-induced vascular injury

In the LSR model, neutrophil infiltration at the LPS-TNFα injection site has been well-documented (32, 33). To identify the primary cell type responsible for vascular inflammation, we selectively depleted neutrophils in wild-type mice using Gr-1 antibodies before administering LPS-TNFα. In neutrophil-depleted mice, no signs of vascular injury were observed, as illustrated by the absence of vascular hemorrhage (Figure 2A). These results confirm that neutrophils are essential mediators of LPS-TNFα-induced vascular injury.

**Figure 2.**
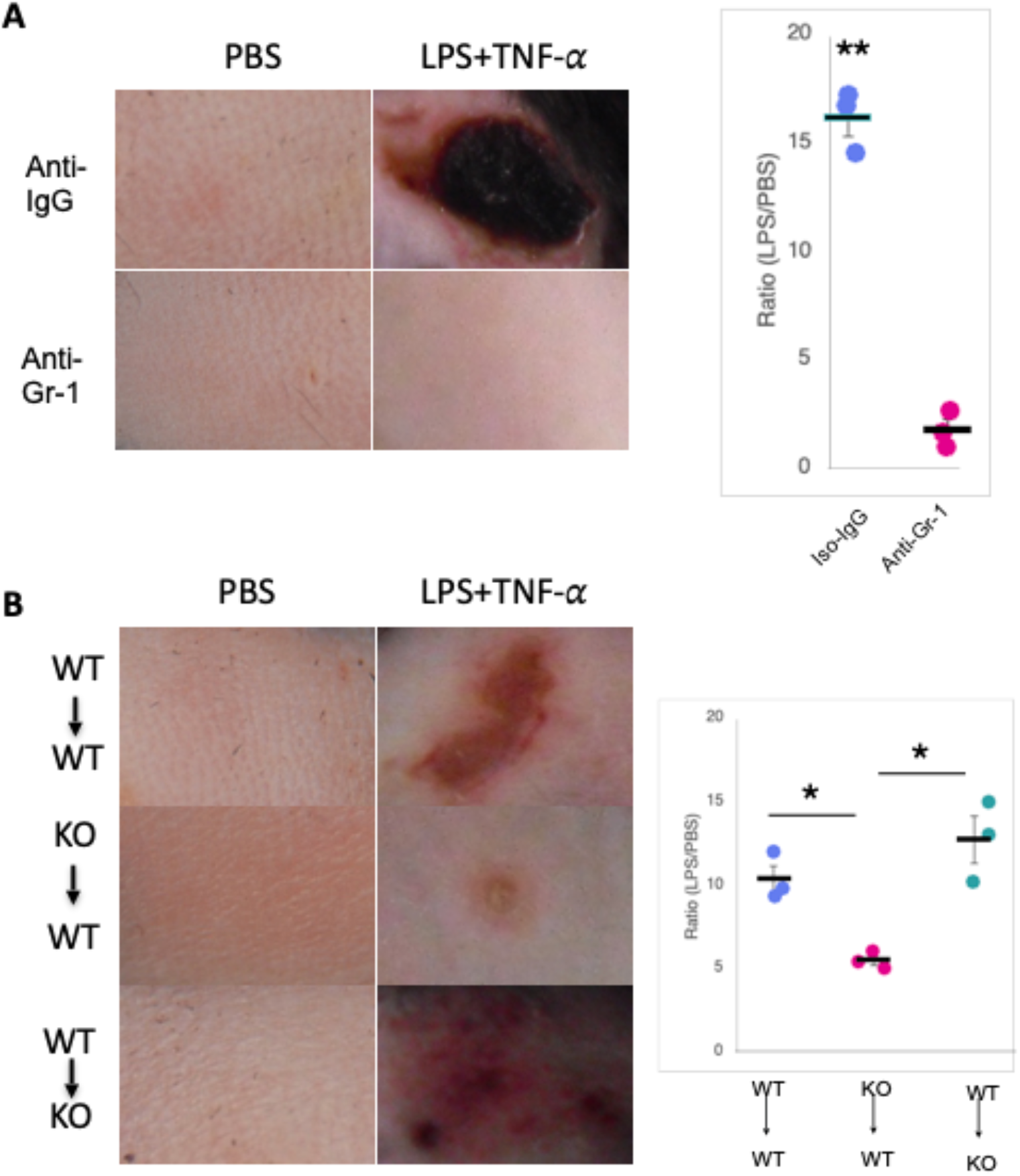
JSP1-dependent neutrophils were responsible for LPS-TNF-α-induced vascular injury. (A) Macroscopic appearance of dorsal skin in WT mice injected subcutaneously with PBS or LPS-TNF-α, which received either anti-IgG or anti-Gr-1 antibody 48 hours before PBS or LPS-TNF-α challenge. (B) Macroscopic appearance of dorsal skin in mice that were intravenously injected with bone marrow neutrophils (2×10^6^) isolated from either *Jsp1*^-/-^ (KO to WT) or WT mice (WT to KO or WT to WT). The degree of hemorrhage in mice was quantified by densitometry analysis of skin samples treated with either PBS or LPS injection. Each dot represents the ratio of the degree of hemorrhage obtained from a pair of mice (LPS–TNF-α– treated over PBS-treated). \**P*<0.05; \*\**P*<0.01, based on three pairs of mice in each group.

To investigate further the role of JSP1 in neutrophil-mediated inflammation, we performed adoptive transfer experiments using neutrophils from different genetic backgrounds. Intravenous transfer of *Jsp1*^*-/-*^ neutrophils into WT mice significantly diminished vascular injury upon LPS-TNFα challenge, suggesting that JSP1-deficient neutrophils have a reduced pro-inflammatory capacity. Conversely, transferring WT neutrophils into *Jsp1*^*-/-*^ mice restored vascular injury to levels comparable to those observed in WT mice (Figure 2B), demonstrating that the presence of JSP1-expressing neutrophils is sufficient to drive inflammation and vascular damage. These observations establish that neutrophils are the mediators of vascular injury in the LSR model and that their pro-inflammatory function is JSP1-dependent. This suggests that JSP1 plays a critical role in neutrophil activation or in signaling pathways that contribute to LPS-TNFα-induced vascular permeability and inflammation.

### JSP1 did not affect neutrophil maturation or CD11b expression

To determine whether JSP1 influences neutrophil maturation or surface marker expression, we performed flow cytometry analysis on bone marrow-derived mature neutrophils from WT and *Jsp1*^−^/^−^ mice. The analysis revealed no significant difference in the quantity of mature neutrophils between the two genotypes, indicating that JSP1 is not required for neutrophil development or homeostasis (Figure 3A).

**Figure 3.**
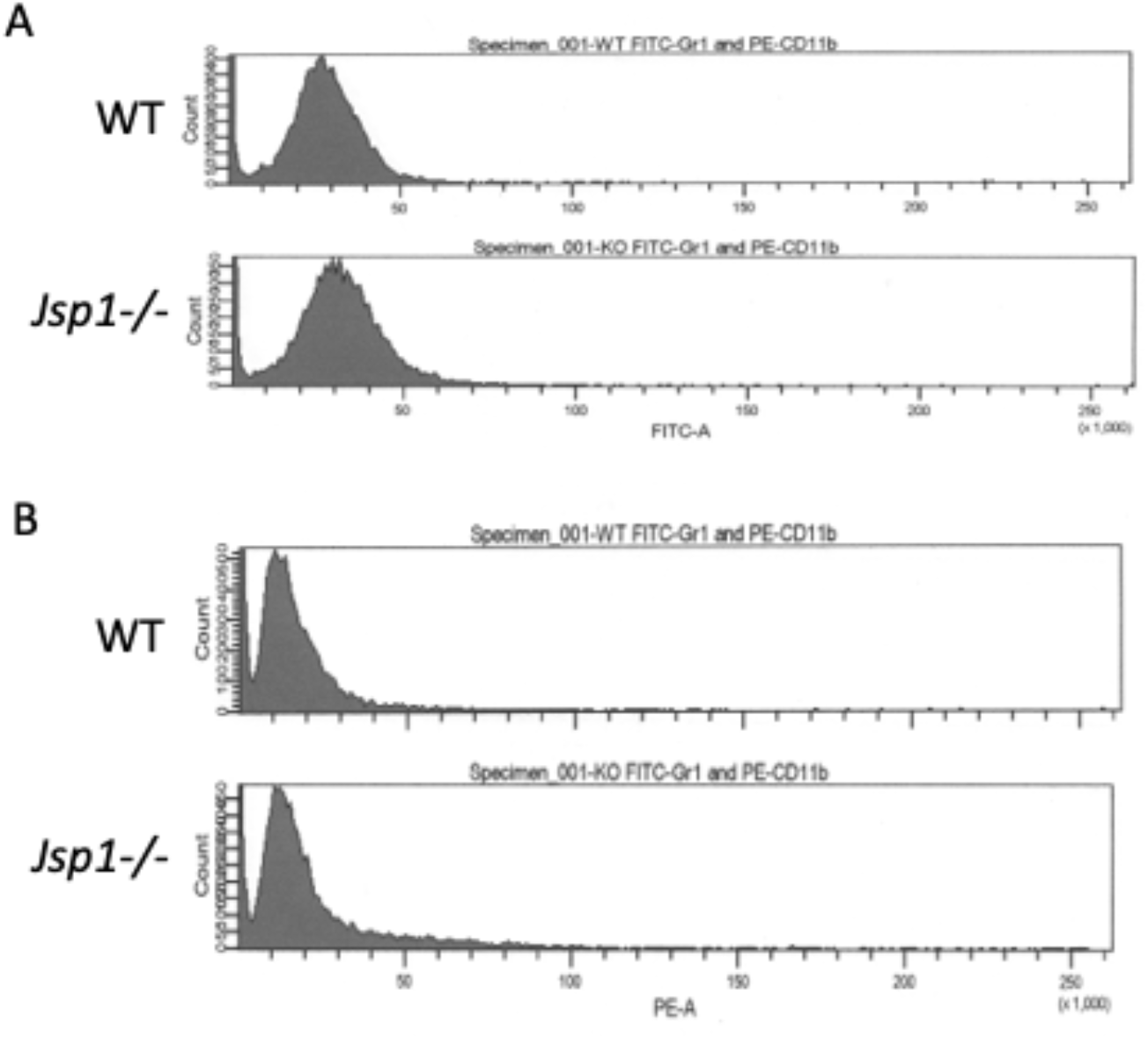
The absence of JSP1 did not alter the quantity of mature neutrophils or the expression of integrin subunit CD11b. Bone-marrow derived neutrophils from either WT or *Jsp1*^-/-^ mice were double immunostained for Gr-1, a marker of mature neutrophils, and integrin subunit CD11b. (A) Flow cytometry analysis to illustrate the surface expression of Gr-1, indicating the quantity of the mature neutrophils. (B) Flow cytometry analysis to illustrate the surface expression of CD11b, indicating the expression levels of the integrin subunit.

Additionally, we assessed the expression of integrin subunit CD11b, also known as integrin alpha M, which is a key surface marker involved in neutrophil adhesion and recruitment. The results showed that CD11b expression levels were comparable between WT and *Jsp1*^−^/^−^ neutrophils (Figure 3B), suggesting that JSP1 does not regulate the surface expression of this integrin.

These findings indicate that the pro-inflammatory role of JSP1 in neutrophil-mediated vascular injury is not due to differences in neutrophil abundance or CD11b expression. Instead, JSP1 likely influences neutrophil function through intracellular signaling mechanisms rather than through changes in neutrophil maturation or integrin expression.

### JSP1 promoted neutrophil adhesion *via* integrin activation

Integrin activation on the neutrophil surface facilitates neutrophil adhesion (33, 34). To examine the role of JSP1 in integrin-mediated neutrophil adhesion, we utilized poly-RGD-coated plates, which serve as integrin cross-linkers to mimic extracellular matrix interactions. Neutrophils from *Jsp1*^−^/^−^ mice exhibited reduced adhesion compared to WT neutrophils (Figure 4), suggesting that JSP1 is required for optimal integrin activation in neutrophils. This observation suggests that JSP1 facilitates neutrophil adhesion by enhancing integrin signaling, which may contribute to its pro-inflammatory role in vascular injury.

**Figure 4.**
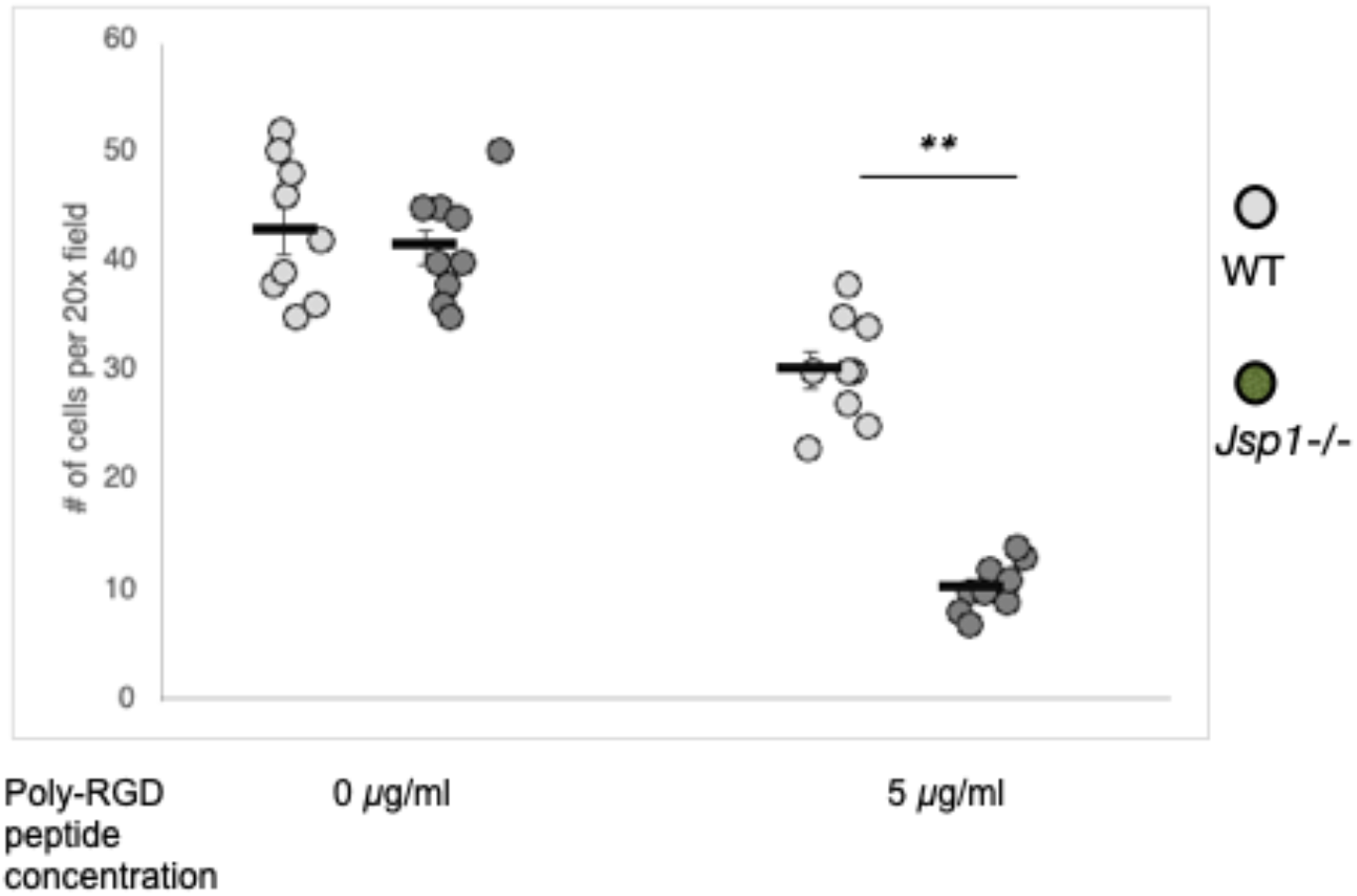
Integrin engagement activates JSP1-dependent neutrophil adhesion. Neutrophils from either WT or *Jsp1*^−^/^−^ mice were plated on poly-RGD–peptide–coated plates to promote integrin engagement. After 15 min of incubation, adherent neutrophils were quantified. Data are presented as mean ± S.E.M. Each dot represents the total number of adherent cells observed under a 20× objective lens. *P* < 0.01, *n* = 9 fields at 200× magnification.

### SRC Family Kinase activation is essential for LPS-TNFα-induced vascular injury

In neutrophils, SRC family kinases HCK, FGR, and LYN, are critical regulators of integrin signaling. Consistently, *Hck*^−^*/*^−^*Fgr*^−^*/*^−^*Lyn*^−^*/*^−^ mice exhibit markedly reduced neutrophil adhesion (34). The spleen tyrosine kinase SYK associates with SRC family kinases specifically in adherent neutrophils bound to fibrinogen (35). Moreover, both HCK and SYK have been implicated as key mediators of neutrophil activation in the LSR mouse model (33). Together, these findings suggest that activation of SRC family kinases is essential for mediating the inflammatory response and vascular damage following LPS–TNFα stimulation.

To test directly the role of SRC family kinases in neutrophil adhesion and vascular injury, we used the SRC tyrosine kinase inhibitor PP2 (36). Treatment of WT neutrophils with PP2 markedly reduced their adhesion to fresh mouse serum–coated plates (Figure 5A). Furthermore, co-injection of PP2 with LPS and TNFα in WT mice resulted in a marked attenuation of vascular injury, as demonstrated by decreased vascular permeability and reduced inflammatory responses (Figure 5B). These observations highlight SRC family kinase activation as critical for neutrophil adhesion and subsequent vascular injury in the LSR mouse model.

**Figure 5.**
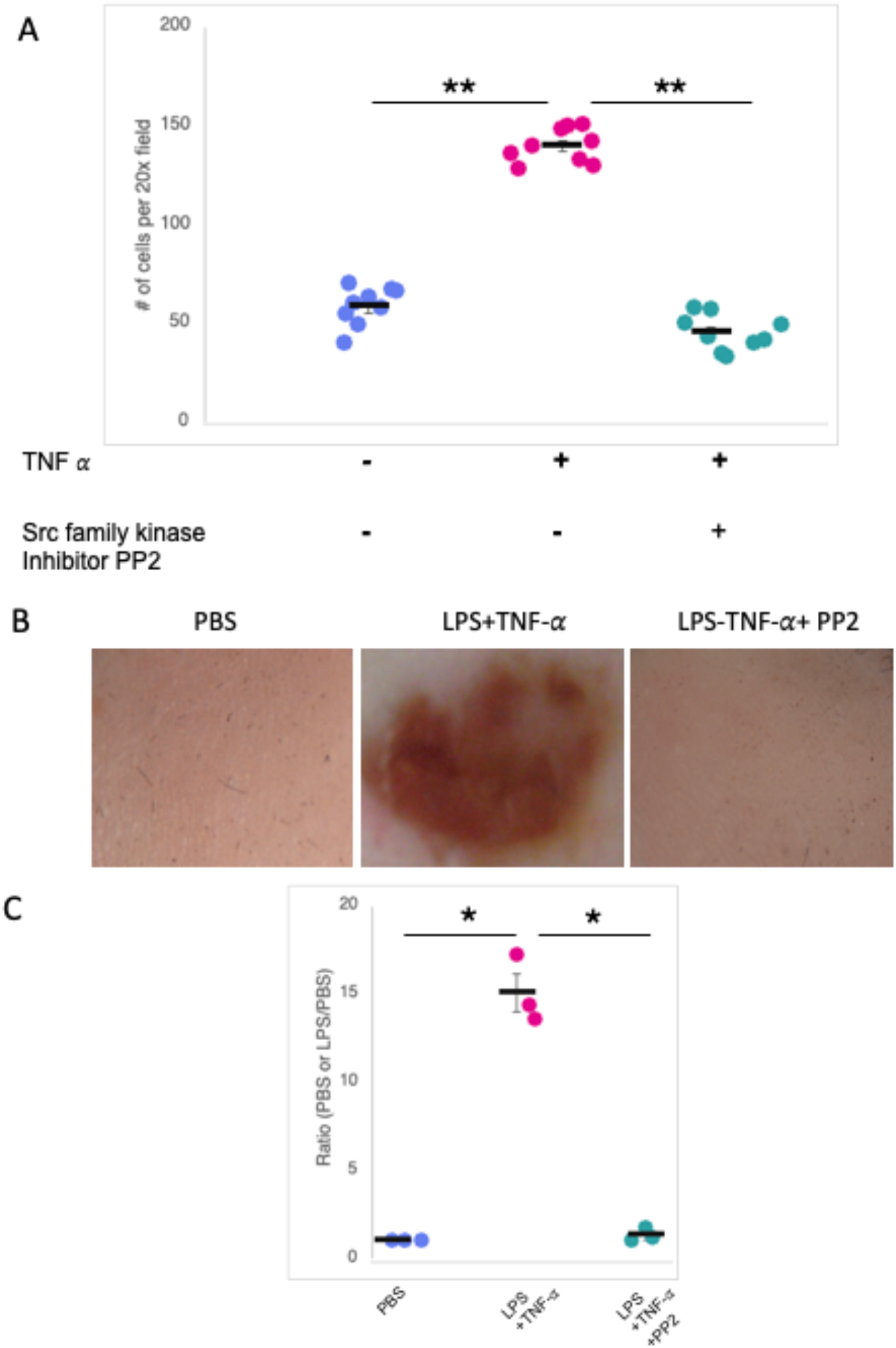
PP2, an inhibitor of SRC family kinases, attenuated neutrophil adhesion and LPS-induced vascular inflammation in WT mice. (A) Quantitative analysis of TNF-α-stimulated WT neutrophil adhesion to fresh mouse serum-coated plates in the presence of SRC family kinase inhibitor PP2. Neutrophils were incubated in plates for 40 minutes, followed by washing and microscopic counting of attached neutrophils across 9 fields at 200x magnification. Each dot represents the total number of adherent cells observed under a 20× objective lens. Data are presented as the mean ± S.E.M. ** *P*<0.01 (B) Morphological examination of dorsal skin in WT mice subcutaneously injected with either PBS, LPS+TNF-α, or LPS+TNF-α along with the SRC tyrosine kinase inhibitor PP2. (C) The degree of hemorrhage in mice was quantified by densitometry analysis of skin samples treated with PBS, LPS+TNF-α, or LPS+TNF-α+PP2 injection. Each dot represents the ratio of the degree of hemorrhage obtained from a pair of mice (Treated over PBS-injected). \**P*<0.05, based on three pairs of mice in each group.

### JSP1 positively regulates SYK and HCK phosphorylation in integrin-activated neutrophils

To investigate the molecular mechanism by which JSP1 modulates neutrophil adhesion and activation, we examined the phosphorylation status of key signaling proteins involved in integrin-mediated responses. SYK, a critical tyrosine kinase in integral signaling, exhibited reduced activation in integrin-stimulated *Jsp1*^−^/^−^ neutrophils compared to WT neutrophils (Figure 6A). Similarly, decreased phosphorylation of HCK was observed in *Jsp1*^−^/^−^ neutrophils upon integrin activation (Figure 6B). Interestingly, and consistent with its function as a protein phosphatase, we observed that JSP1 was able to dephosphorylate the C-terminal inhibitory phosphorylation site in HCK (Figure 6C).

**Figure 6.**
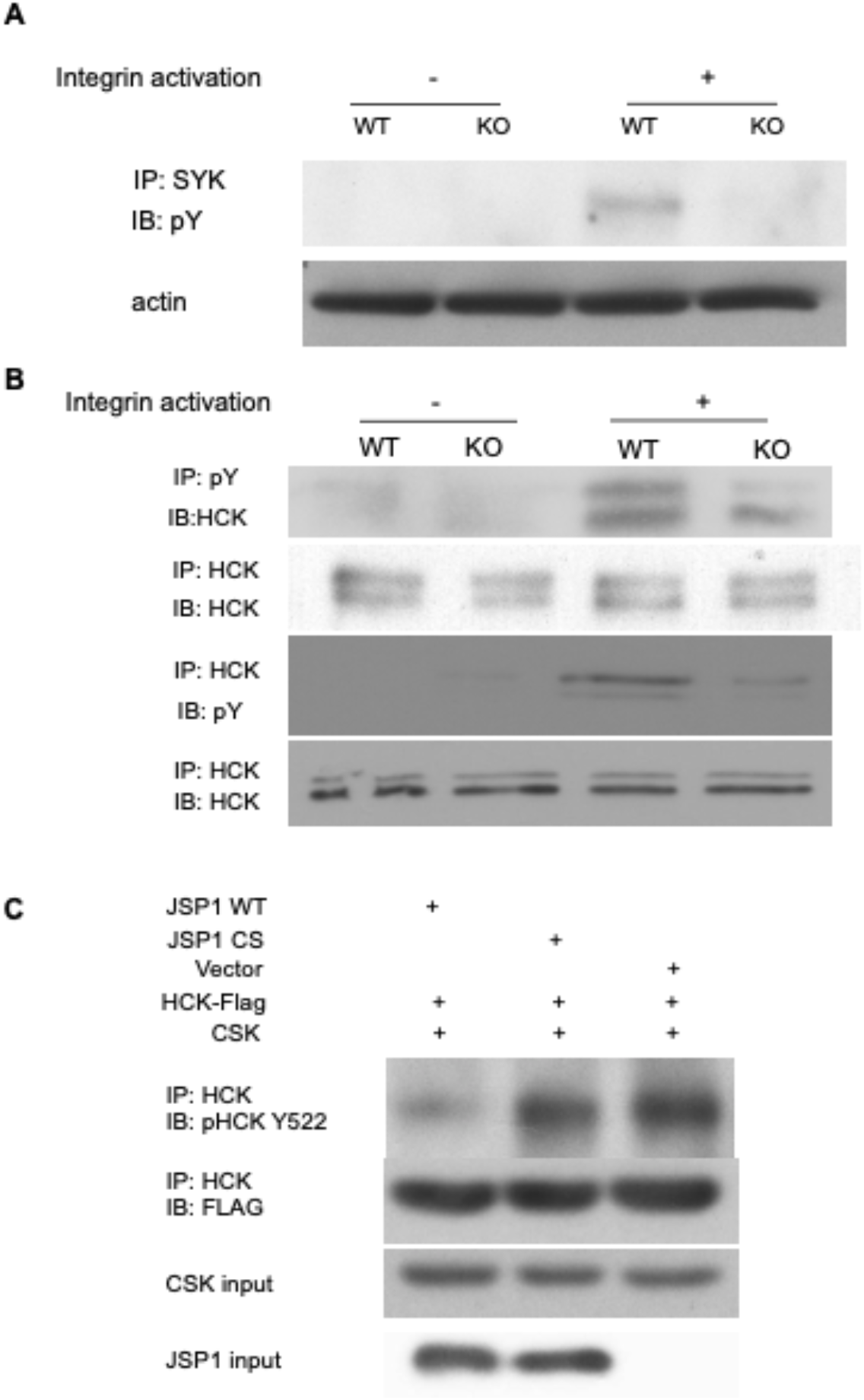
Tyrosine phosphorylation of SYK and HCK is down-regulated in integrin-activated *Jsp1*^-/-^ neutrophils. (A) Lysates of wild type (WT) or *Jsp1*^-/-^ (KO) neutrophils, either plated on a poly-RGD-coated surface (+ integrin activation) or suspended (-integrin activation) in assay media, were prepared after 15-minute incubation and immunoprecipitated with an anti-SYK antibody. The resulting immunoprecipitates were analyzed to assess the level of SYK tyrosine phosphorylation with anti-phosphotyrosine (4G10) antibody. (B) Lysates of neutrophils treated as described in (A) were also immunoprecipitated with either an anti-phosphotyrosine (4G10) antibody or an anti-HCK antibody to evaluate the tyrosine phosphorylation of HCK and overall tyrosine phosphorylation levels. (C) JSP1 activates HCK by dephosphorylating the negative regulatory tyrosine 522. Lysates of 293T cells transfected with either JSP1 WT, JSP1 CS (inactive mutant), or vector control, along with flag-tagged HCK and CSK, were immunoprecipitated with an anti-HCK antibody. The immunoprecipitates were then analyzed by immunoblotting with anti-pHCK Y522 to detect phosphorylation at tyrosine 522, and with anti-FLAG to confirm the expression of the transfected constructs.

Overall, these observations are consistent with an important role of JSP1 in fine-tuning neutrophil activation by upregulating SYK and HCK phosphorylation, thereby influencing integrin-mediated adhesion and inflammatory responses.

## Discussion

Members of the PTP family are critical regulators of signaling pathways through their ability to dephosphorylate diverse substrates. Within that family, classical DUSPs, such as those targeting MAPKs, typically function as negative regulators that can influence the intensity and duration of signaling by dephosphorylating and inactivating ERK, JNK, or p38 (1-6). Conversely, protein phosphatases can also function to switch on signaling. For example, cdc25 acts positively to promote cell-cycle progression by dephosphorylating inhibitory residues on cyclin-dependent kinases (CDKs) (37). These contrasting activities highlight the dual potential of phosphatases to either dampen or amplify signaling events depending on the cellular context and substrate specificity. In this study, we highlight a further example of a phosphatase that may act positively to promote a signaling response.

Since its identification as a unique DUSP capable of activating JNK signaling (7,8), JSP1 (DUSP22) has garnered attention as a regulator of diverse physiological and pathophysiological processes, including immunity and cancer. Structural studies revealing a shallow and accessible catalytic cleft, consistent with its capacity to act on multiple substrates and pathways (38). Our findings extend these observations by demonstrating that JSP1 also plays a pivotal role in neutrophil adhesion and inflammatory vascular injury.

The Local Shwartzman Reaction (LSR) is an animal model of the innate immune response, including JNK-driven vascular inflammation (30, 31). This is a two-stage model. Firstly, a priming injection of endotoxin (LPS) sensitizes the innate immune system by recruiting inflammatory cells, such as neutrophils, to the injection site. A second injection of TNFα, or endotoxin to activate release of inflammatory cytokines such as TNFα, triggers an acute inflammatory response in which initiation of JNK signaling cascades is of particular importance (31, 39). In this study, we have established that systemic deletion of JSP1 protects against LPS/TNFα-induced vascular injury in this model. This protection correlates with impaired neutrophil adhesion and reduced activation of integrin signaling, implicating JSP1 as a key upstream regulator in the inflammatory cascade. Our adoptive transfer experiments reveal a cell-intrinsic requirement for JSP1 in neutrophils, underscoring the fact that its pro-inflammatory role is directly attributable to neutrophil function and is not secondary to systemic alterations. Mechanistically, neutrophils lacking JSP1 displayed diminished phosphorylation of HCK and SYK—critical kinases in the integrin signaling pathway. These data suggest that JSP1 facilitates integrin-mediated signaling in neutrophils, possibly enabling the assembly or stabilization of signaling complexes at adhesion sites and strengthening the notion that JSP1 operates as a cell-specific regulator of innate immunity.

The JNK signaling pathway can be activated in response to a wide array of stimuli from cytokines to tissue damage. In turn, this underlies fundamental aspects of the inflammatory response, from recruitment and activation of immune cells to engaging the adaptive immune system. Disruption of such signaling has been implicated in a range of major diseases, including vascular diseases. The development of approaches to target JNK activation in such contexts holds major therapeutic potential. By expanding the functional repertoire of JSP1 to the modulation of neutrophil signaling and vascular inflammation, our study illustrates a new potential mechanism to achieve this. The identification of JSP1 as a positive regulator of SRC family kinase activation in neutrophils provides a mechanistic link between phosphatase activity, integrin signaling, and vascular injury. Importantly, these insights position JSP1 as a potential therapeutic target for conditions characterized by excessive neutrophil activation and vascular damage, including sepsis, autoimmune vasculitis, and ischemia-reperfusion injury. Future studies will be needed to determine the precise molecular mechanism by which JSP1 promotes HCK and SYK activation, balancing direct dephosphorylation of inhibitory residues in the SRC family kinases with indirect effects on scaffolding functions, and to define potential links between SRC and JNK activation in this system (40). Furthermore, there have been reports of the identification of two compound inhibitors of JSP1 activity (18, 41); it will be important to assess whether selective inhibition of JSP1 with small-molecule drug candidates can attenuate neutrophil-driven pathology in clinically relevant models.

In conclusion, our findings reveal a previously unrecognized role of JSP1 in neutrophil adhesion and vascular inflammation through the regulation of SRC family kinase signaling. By bridging phosphatase activity with innate immune function, this work highlights JSP1 as a novel and promising target for therapeutic modulation of inflammatory vascular injury.

## Methods and Materials

### Materials

Reagents and antibodies are summarized in Table 1.

**Table 1.** Reagents and Antibodies

| **Reagent/Antibody** | **Source** | **Product/Catalog No.** |
| --- | --- | --- |
| LPS | Sigma-Aldrich | L6529, 1mg |
| TNFα | Sigma-Aldrich | T5944, 10 μg |
| PP2 | Sigma-Aldrich | 529573 |
| 10xHBSS, Ca<sup>2+</sup>/Mg<sup>2+</sup>-free | Thermo Fisher Scientific | 14185052 |
| 100% Percoll | Sigma-Aldrich | P1644 |
| Nyco-prep | Accurate Chemical & Scientific Corp. | AN1106865 |
| RBC lysis buffer | Sigma-Aldrich | R7757 |
| poly-RGD peptides | Sigma-Aldrich | F5147- 1 mg |
| Protein G agarose bead slurry | Cell Signaling | 37478 |
| pSYK (Y323) antibody | Cell Signaling | 2715 |
| HCK antibody | Santa Cruz Biotechnology | sc-1428 |
| 4G10 antibody | Millipore | 05-321 |
| pHCK (Y522) antibody | Thermo Fisher Scientific | 44-912 |

### Mice

The mouse studies were performed in accordance with procedures approved by the IACUC at CSHL and NIH Guide for Care and Use of Laboratory Animals.

### JSP1 knockout mice generation

Two-to three-month-old mice were used. C57BL/6 mice were purchased from Taconic Farms Inc., NY. JSP1 knockout (*Jsp*1^-/-^) mice were generated by targeted deletion of Exon III of the JSP1 gene and backcrossed at least ten times to the C57BL/6 background (CEPTYR Inc.)

### Mouse model of local Schwartzman reaction (LSR)

The physiological functions of JSP1 were explored in the mouse model of Local Shwartzman Reaction (LSR, 30-32).

### Anesthesia and mouse preparation for injection

In brief, on the first day of injection, mice were marked and anesthetized in an isoflurane chamber. Anesthetized mice were rubbed with Nair Hair Removal Cream on their dorsal hair and left in the isoflurane chamber with padding for at least 5 minutes. Then the creamed hair was removed by a soapy warm water-immersed gauze followed by a regular tap water-immersed one. The exposed skin was sterilized by 70% ethanol.

### Subcutaneous injection of LPS and TNFα

On Day 1, mice were injected subcutaneously on the dorsal skin with 80 µl of either sterile PBS or LPS (1 mg/ml). On Day 2, 80 µl of either sterile PBS or TNFα (0.2 µg) was administered subcutaneously at the same dorsal site as the Day 1 injection.

### Macroscopic examination of skin lesion was done on Day 3

Mice were anethestized in an isoflurane chamber with padding and macroscopically examined for vascular inflammation. Pictures were taken with a Canon digital camera. The relative levels of hemorrhage were quantified by densitometry of macroscopic images using ImageJ software (National Institutes of Health).

### Neutrophil isolation

Preparation of bone-marrow-derived mouse neutrophils was described previously (32). In brief, mouse bone marrow cells flushed from femurs and tibias were resuspended in 3 ml Ca^2+^/Mg^2+^-free Hank’s balanced salt solution (HBSS) containing 1% BSA, which were loaded onto a discontinuous density gradient containing 3 ml of a 72% Percoll under 3 ml of Nyco-prep. After centrifugation at 1000 x g for 20 minutes (without brake when slowing down) at room temperature, cells at the interface between the 72% Percale and Nyco-prep were collected and washed with Ca^2+^/Mg^2+^-free HBSS. The cell pellet was resuspended in 1 ml of RBC lysis buffer and gently mixed for 1 minute. The cell mixture was diluted with 15 ml of HBSS, pelleted after centrifugation at 500 x g for 7 minutes and resuspended in 500 µl of 1xHBSS /1% BSA. Cells were quantitatively estimated using a hemacytometer.

### Neutrophil Adhesion Assay

6 cm tissue culture dishes were coated by incubating with either 1.5 ml of fresh mouse serum or various concentrations of poly-RGD peptides in a humidified incubator at 37ºC for 1.5 hours. The coated plates were then washed twice with 2 ml of 1x HBSS/Ca^2^+/Mg^2^+/20 mM HEPES and resuspended in 1x HBSS/1% BSA to remove unbound peptides or serum components.

Freshly isolated bone marrow-derived neutrophils from either wild-type (WT) or *jsp1*-/-mice were prepared. Half of the neutrophils were kept on ice as the “suspension” fraction, while the other half were plated onto either poly-RGD-coated plates and incubated at 37ºC for 15 minutes or fresh mouse serum-coated plates and incubated for 40 minutes to allow adhesion. After incubation, the non-adherent cells were removed by carefully aspirating the supernatant, and the adherent cells were lysed directly on the plate with 0.5 ml of lysis buffer.

### Protein Extraction and Immunoprecipitation

For immunoprecipitation (IP) experiments, cells were lysed in RIPA buffer (25 mM Tris-HCl, pH 7.4, 150 mM NaCl, 1% NP-40, 1% sodium deoxycholate and 0.1% SDS) supplemented with protease inhibitor cocktail. For immunoblotting, cells were lysed in NP-40 lysis buffer (LB). Lysis was performed in a cold room with gentle rocking for 30 minutes. Lysates were then centrifuged at 13,000 rpm for 15 minutes, and the supernatant was collected. Protein concentration was determined using the Bradford protein assay.

For IPs, 1 mg or 750 µg of lysate was incubated with 20 µl of Protein G agarose bead slurry overnight at 4ºC with continuous rocking. The next day, 10 µl of specific antibodies were added to the lysate-bead mixture and incubated at 4ºC for 90 minutes with rocking.

Immunoprecipitates were collected by centrifugation and washed three times with LB. Proteins were eluted by boiling with 2X SDS-PAGE loading buffer followed by immunoblotting analysis using the appropriate antibodies, including pSYK (Y323), HCK, pTyr 4G10 and pHCK (Y522).

### Statistical Analysis

All data are expressed as means ± standard error of the mean (SEM). An unpaired Student’s *t*-test was used to compare two groups for all experiments. Significance was set at *P*<0.05.

